# Local changes dominate variation in biotic homogenization and differentiation

**DOI:** 10.1101/2022.07.05.498812

**Authors:** Shane A. Blowes, Brian McGill, Viviana Brambilla, Cher F. Y. Chow, Thore Engel, Ada Fontrodona-Eslava, Inês S. Martins, Daniel McGlinn, Faye Moyes, Alban Sagouis, Hideyasu Shimadzu, Roel van Klink, Wu-Bing Xu, Nicholas J. Gotelli, Anne Magurran, Maria Dornelas, Jonathan M. Chase

## Abstract

It is commonly thought that the biodiversity crisis includes widespread decreases in the uniqueness of different sites in a landscape (biotic homogenization). Using a typology relating homogenization and differentiation to local and regional diversity changes, we synthesize patterns across 283 metacommunities surveyed for 10-91 years, and 54 species checklists (13-500+ years). On average, there is a 0.2% increase in species shared among communities/year (i.e., weak homogenization), but across data sets, differentiation frequently occurs, with no statistically significant change being most common. Local (not regional) diversity frequently underlies composition change, and homogenization is strongly associated with checklist data that have longer durations and large spatial scales. Conservation and management can benefit from the multiscale perspective used here as it disentangles the implications of both the differentiation and homogenization currently unfolding.

**One-Sentence Summary:** Biotic homogenization is most prevalent at large temporal and spatial scales.

---

Humans are fundamentally altering the Earth’s land cover, climate, water and nutrient cycles (*1*). With this ever growing human footprint, Earth’s biodiversity is inevitably changing (*2*). There is substantial evidence that humans are accelerating the global extinction rate (*3*). However, at local scales, no overall directional trend, or slight increases in diversity are typically detected amidst substantial variability (*4–7*), though not without controversy (*8*). One common explanation for the perceived discrepancy between declining diversity globally and unchanged diversity at local sites on average, is biotic homogenization (*9–11*).

Biotic homogenization is a frequently observed pattern by which spatially distinct locations become more similar to one another in species composition through time (*12–14*). Two opposing forces can lead to homogenization: increased numbers of widespread (high occupancy) species (e.g., often non-native species), or extirpation of rare (low occupancy) species from regions. One driver for homogenization is when anthropogenic use creates environments with high habitat similarity (e.g., agricultural practices or harvesting). However, human activities can also create more heterogeneous environments (*15*), for example, by habitat fragmentation and other processes, which can lead to biotic differentiation. Moreover, non-native species that are introduced but do not become widespread, or when formerly widespread species are locally extirpated can also lead to differentiation (*16–17*). While both biotic homogenization and differentiation are frequently observed and theoretically expected under different scenarios, an empirical assessment of their prevalence in surveys of biodiversity change through time is lacking. Moreover, despite the intrinsic connection between homogenization and differentiation of community composition and rates of change in diversity across spatial scales (*16–17*), the empirical relationships among these have not been well quantified and synthesized.

## Visualizing Whittaker’s scale-explicit diversity partition

We can illustrate the relationship between homogenization and differentiation and rates of change across spatial scales using Whittaker’s (*18*) diversity partition, where the diversity of a single site (i.e., a local site) is *α*-diversity, and the combined diversity of several local sites (i.e., a region) is *γ*-diversity. Variation in species composition among local sites, referred to as *β*-diversity, is given by: *β* = *γ*/ᾱ (where ᾱ is the average local diversity across sites in a region). Therefore, if rates of change in *α*- and *γ*-diversity through time are not equal, *β*-diversity changes through time. Moreover, these changes in *β*-diversity can be mathematically linked to changes in the number of sites species occupy (i.e., occupancy). Average occupancy, or the proportion of sites within a region occupied by species *i* (o*_i_*), is related to Whittaker’s formula by *β* = *γ*/ᾱ = *γ*/(*Σo_i_*) = *γ*/(*γ*ō) = 1/ō (*19*). Thus, whether β-diversity decreases through time (homogenization) or increases through time (differentiation) is directly linked to change in diversity at two different scales, and to changes in the average proportion of sites that species occupy (Fig. 1).

**Figure 1:**
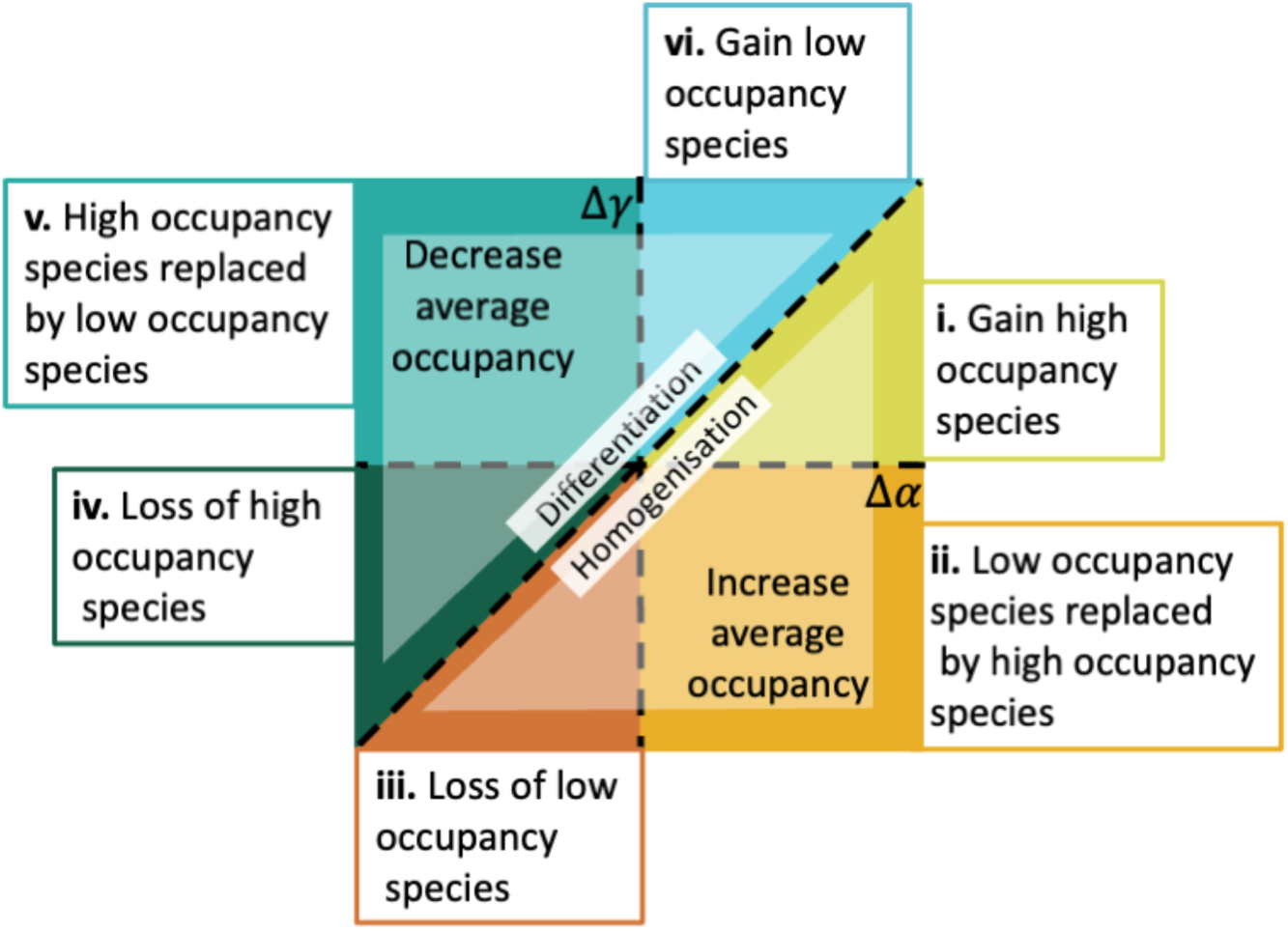
Mechanisms that underpin changes in spatial variation of species composition can be understood by examining the relationship between changes in regional- and local-scale species richness through time. When richness changes at regional (Δ*γ*) and local (Δ*α*) scales are calculated as proportional changes, assemblages below the dashed 1:1 line, i.e., Δ*γ* < Δ*α*, are being homogenized, *β*-diversity is decreasing, and average occupancy is increasing. Conversely, assemblages above the diagonal dashed 1:1 line, i.e., Δ*γ* > Δ*α*, are differentiating, *β*-diversity is increasing, and average occupancy is decreasing. In regions where (i) Δ*γ* < Δ*α*, and Δ*γ* >0, Δ*α* > 0, the number of species with high occupancy (i.e., that occupy the majority of sites in the region) is increasing; (ii) Δ*γ* < Δ*α*, Δ*γ* < 0 and Δ*α* > 0, average occupancy is increasing, e.g., due to species with low occupancy being replaced by those with high occupancy; (iii) Δ*γ* < Δ*α* and Δ*γ*, Δ*α* < 0, the number of species with low occupancy is decreasing; (iv) Δ*γ* > Δ*α* and Δ*γ*, Δ*α* < 0, the number of species with high occupancy is decreasing; (v) Δ*γ* > Δ*α*, Δ*γ* > 0 and Δ*α* < 0, species with low occupancy (i.e., occupy few sites in the region) are replacing those with high occupancy; (vi) Δ*γ* > Δ*α* and Δ*γ*, Δ*α* > 0, the number of species with low occupancy (i.e., occupy relatively few sites in the region) is increasing.

A more ecologically informed understanding of scale-dependent biodiversity change can be gained by explicitly considering change in *α*-, *β* - and *γ*-diversity simultaneously. We illustrate this more detailed picture of scale-dependent change in Figure 1, showing six qualitatively distinct scenarios that emerge in the intersecting space of temporal changes in *α*- and *γ*-diversity. The 1:1 line (i.e., Δ*γ* = Δ*α*, *Δβ* = 0) represents equal log-proportional changes at both scales (i.e., log *γ*/year = log *α*/year), and delineates the boundary between homogenization and differentiation (Fig.1). When Δ*α*>Δ*γ* (i.e., below the 1:1 line), homogenization occurs due to increased average occupancy; three of the six scenarios are possible. (1) Increased numbers of high occupancy species, due for example, to environmental changes that favor widespread, generalist and/or non-native species, could drive increases in *α*-diversity proportionately more than *γ*-diversity, leading to homogenization (with Δ*α*>Δ*γ*>0, Fig. 1i). (2) If widespread species replace low occupancy species, average occupancy increases, and homogenization is associated with diversity gains at the *α*-scale and losses at the *γ*-scale (Δ*α*>0, Δ*γ*<0, Fig. 1ii). (3) If low occupancy species (e.g., endemic species restricted to few sites), are regionally extirpated, and homogenization is associated with diversity loss at both the *α*- and *γ*-scale (Δ*γ* <Δ*α* <0, Fig. 1iii). These distinct scenarios all describe cases of biotic homogenization, but variation in the nature of scale-dependent change could have different implications for policy and conservation. Three further parallel and distinct scenarios of differentiation (and decreasing average occupancy) through time are also possible. (4) Fewer widespread or high occupancy species could, for example, result from increased habitat heterogeneity, and greater losses at *α*- relative to *γ*-scales combine to increase *β*-diversity (Δ*α*<Δ*γ*<0, Fig. 1iv). Increased habitat heterogeneity could also lead to increases in *γ*-diversity, accompanied by either: (5) *α*-diversity declines if low occupancy species replace high occupancy species (Δ*α*<Δ*γ*, Δ*α*<0, Δ*γ*>0, Fig. 1v); or, (6) increased *α*-diversity due to increased numbers of low occupancy species (Δ*γ*>Δ*α*>0, Fig. 1vi; see Fig. S1 for further illustration of scale-dependent variation).

To assess overarching trends in scale-dependent biodiversity change, we use a compilation of 337 datasets documenting taxonomic diversity through time across spatial scales (Fig. S2) and the typology of Figure 1 to estimate changes in *α*-, *β*-, and *γ*-diversity. Among these biodiversity datasets, there were 283 studies where we could calculate sample effort-controlled species richness from local sites (*α*-diversity) and from the broader region (*γ*-diversity), for which we set a minimum of at least four sites per region and at least ten years between the first and last samples; 173 studies came from already compiled databases (*20–22*), and 110 datasets had a similar structure, but were specifically compiled for this and related studies (see Material and Methods). The remaining 54 datasets were compiled from presence-absence species ‘checklists’ collected from multiple sites (at least four) and time periods separated by a minimum of 10 years; these data typically encompass much longer timespans, documenting historical species composition prior to major human disruption and a more contemporary time period, but only have two time points. We include species checklist data despite their coarse nature as they have made key contributions to our understanding of long-term trends in introductions and extinctions (*23, 24*), as well biotic homogenization (*25, 26*).

We first asked whether there were any general tendencies among all 337 datasets with respect to the six qualitatively distinct scale-dependent biodiversity change scenarios overviewed in Figure 1 (plus the case where there is no change at either scale). We calculated temporal changes in diversity at the smaller, *α*-scale, at which one sample was taken, and a larger *γ*-scale, where diversity was taken as the sum of species in all of the local samples combined. The grain of the *α*-scale and extent of *γ*-scales varied among datasets, ranging, for example, from quadrat samples of plant communities collected over small spatial extents (<< 1km^2^) to species checklists of birds on islands distributed across several oceans. To these data, we fit multilevel models separately to rates of change for each scale calculated as a log-ratio using two time points (start and end of the time series) standardized by the duration between samples (i.e., log[(S_t2_/S_t1_)/(t2 – t1 + 1)], where S is species richness, and t1 and t2 denote the year of first and last sample, respectively); both models adjusted for residual heterogeneity associated with duration and the different data sources (i.e., resurveys and checklists; see Material and Methods).

### Scale-dependent diversity changes and biotic homogenization and differentiation

All possible outcomes were observed across the different data sets (Fig. 2a). In combination, α- and γ-scale changes result in many data sets having no trend in *β*-diversity, many with trends towards homogenization (lower *β*-diversity through time), and many with trends towards differentiation (higher *β*-diversity through time), though few individual data sets had values of *Δβ* that statistically differed from zero (Fig. 2e). Overall, we found a weak mean trend towards homogenization (*Δβ* = -0.002; Fig. 2b). This trend equates approximately to 2 out of 1000 entirely distinct (i.e., no shared species) communities (*27*) being removed per year (rate of change calculated as the distance from the 1:1 line; 90% credible interval: -0.003, -0.0008). Stated differently, this represents an average increase by 0.2%/year of the number of shared species among localities within a region. This mean result was driven by approximately half of the datasets that had only two time points (i.e., checklists [n = 54], and resurvey data [n = 118] where contemporary samples were collected to document changes compared to a historical sample; Fig. S3); in contrast, data with a time series (i.e., multiple years) actually showed a slight tendency to differentiation (mean *Δβ*=0.0002, Fig. S3) rather than homogenization. Homogenization in the data with only two time points was strongly associated with checklist data (Fig. 2a), and residual variation was a decreasing function of sampling duration, especially for checklist samples at the α-scale (Fig. S4). Importantly, we found that homogenization is more likely at intermediate to large spatial and temporal scales, and associated with small effect-sizes for the long duration checklists (Fig. S5). These patterns suggest both differentiation and homogenization are frequent at smaller to intermediate scales as environmental heterogeneity can increase or decrease through time, but as spatial scale increases, homogenization appears to become more likely (*9, 17*).

**Figure 2:**
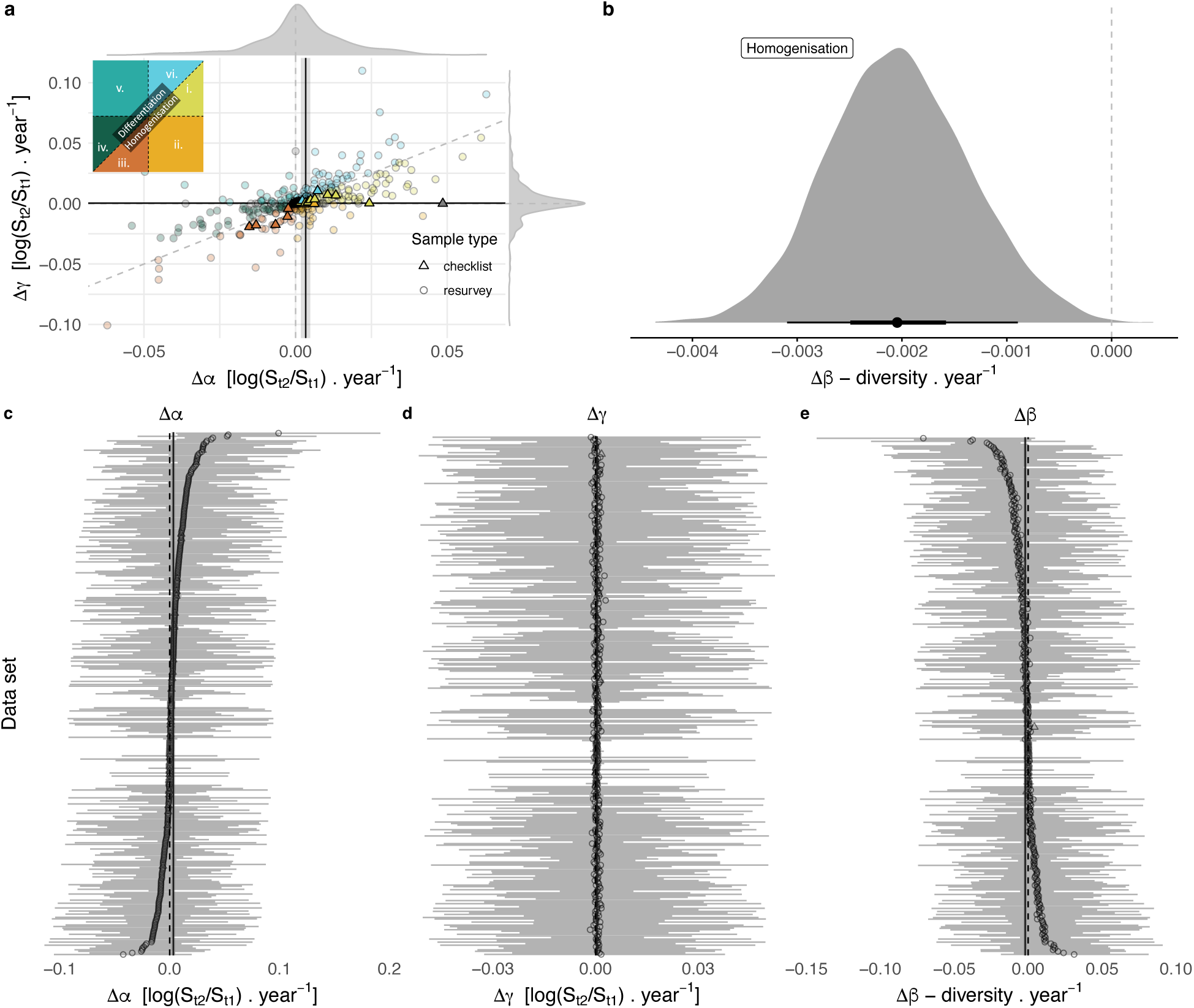
Patterns of homogenization and differentiation across 283 metacommunites and 54 checklists. (a) Empirical estimates of *γ*-scale changes as a function of *α*-scale changes, both axes show log-ratios standardized by the number of years between the estimates, and external kernels show density of empirical data; black lines show median *γ*- and *α*-scale changes, and shading (not visible for *γ*-scale) shows the 90% credible interval from multi-level models fit separately to these data at each scale. Dashed grey lines show x = 0, y = 0, and x = y. (b) Kernel density plot of change in *β*-diversity per year calculated as the distance from 1:1 line (left = homogenization, right = differentiation) of 1000 draws of the posterior distributions of *α*- and *γ*-scale intercepts; black point shows median, bar represents 50% (thick) and 90% (thin) credible intervals. Model estimates of change for each region at the (c) *α*-, (d) *γ*-, and (e) *β*-scales; each point represents a single region, with the bar showing the 90% credible interval; regions are in the same order on panels c-e, arranged by the magnitude of the *α*-scale estimate. Two regions with *Δ*α > 0.1 (0.1 and 0.2) were removed for clarity from panel a; point color represents categories of change from Fig. 1, points falling on boundaries between categories are gray.

Contrary to expectations, the weak trend towards homogenization is associated with gains in α-diversity, more often than losses in γ-diversity. Our conceptual typology (Fig. 1) allows us to parse underlying scale-dependent changes across the different data sets. Most noticeably, variation in β-diversity is primarily driven by changes in α-diversity (Fig. 2c-e). When α-diversity increases, β-diversity goes down since changes in γ-diversity are comparatively small, and vice versa. Thus, the weak overall trend to homogenization (*Δβ* < 0) is associated with a small net average increase in α-diversity (*Δ*α = 0.003; 90% credible interval: 0.002-0.005). This runs contrary to the narrative that no change (or increases) in species richness at smaller spatial scales are due to biotic homogenization and decreased biodiversity at larger spatial scales (*9–11*). Instead, we see smaller positive changes (relative to the smaller α-scale changes) in γ-scale richness (i.e., *Δ*γ <*Δ*α, *Δ*γ = 0.0004, 90% credible interval: 0.0002 - 0.0006) and biotic homogenization associated with increased average occupancy. Across all datasets, we find that increased numbers of widespread, high occupancy species are a key driver of biotic homogenization (see Fig. 1i, Fig. S5, S7).

The subset of data for which time series of species’ relative abundances were available (n = 165; duration range = 10-90 years, median = 16) also show that variation in homogenization and differentiation is frequently dominated by α-scale changes (Fig. 3a, b). These time series data do not include checklists and many of the resurvey studies that had long durations and large spatial scales, and the increased power to detect temporal changes (especially at the γ-scale) revealed more cases where α- and γ-scale changes were approximately equal, and some cases where *Δ*γ > *Δ*α (Fig. 3a, b). This results in overall average α- and γ-scale changes being more closely balanced (Fig. 3c), and average *Δ*β slightly increasing (differentiating), but broadly overlapping zero (Fig. 3d).

**Figure 3:**
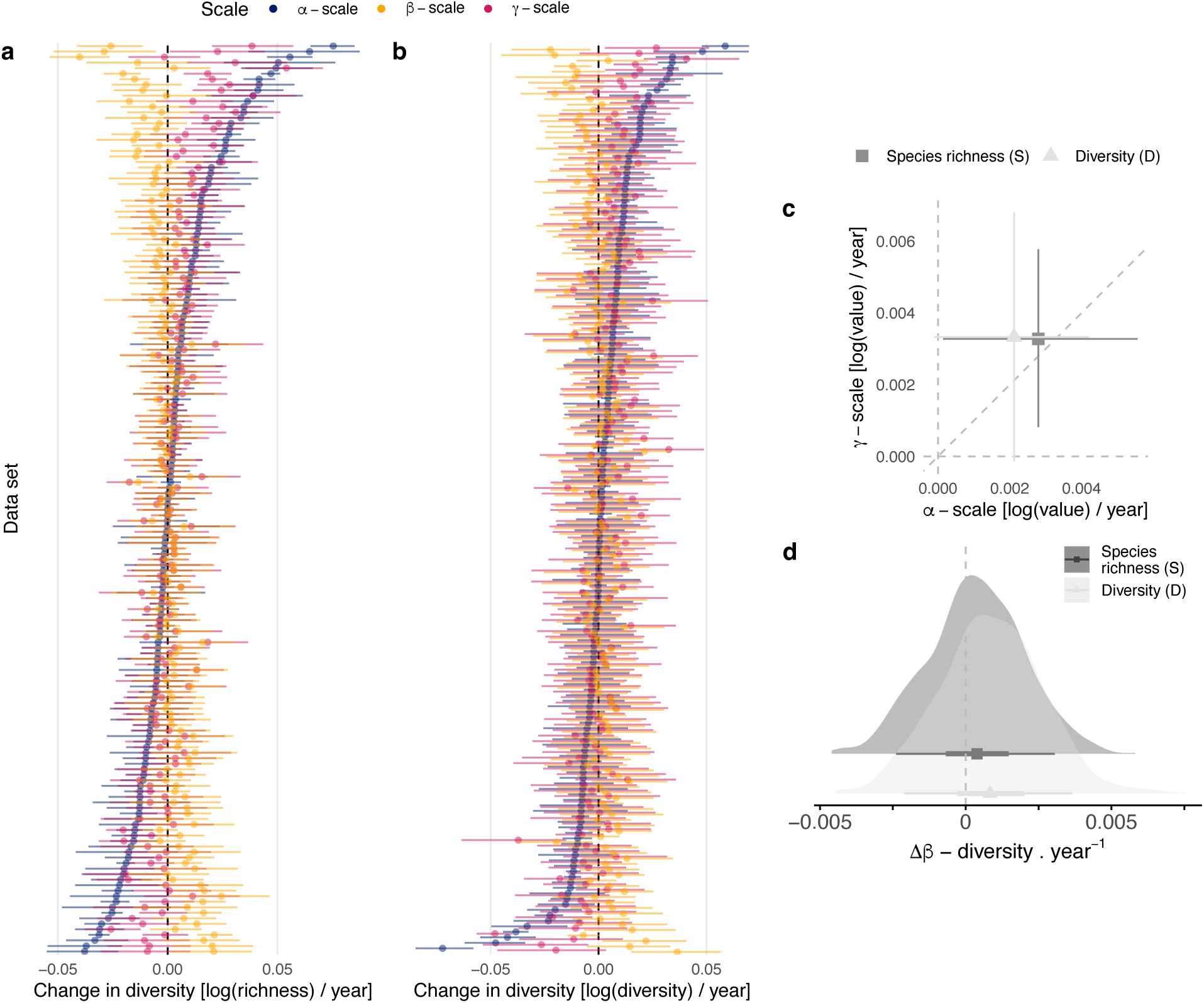
Scale-dependent diversity changes across a subset of 165 metacommunities estimated using models fit to time series of relative abundances. Regional variation in *α*-, β- and *γ*-diversity change estimated using (a) species richness (S), (b) a diversity metric (D), equal to the inverse of Simpson’s concentration (28), and more influenced by numerically abundant species; (c) overall average rates of *α*- and *γ*-scale diversity change, and (d) overall estimates of *Δ*β-diversity. Data sets (regions) in a and b are arranged in order of the magnitude of *α* - diversity changes for each diversity order (i.e., ordered differently on panel a and b). *Δ*β-diversity was estimated as the distance from the 1:1 line and calculated using 1000 draws from posterior distributions, where each x,y coordinate was a single draw from the population-level (fixed effect) slope parameter of *α*- and *γ*-scale models. Points and whiskers show median and 90% credible intervals on a-c, and median, 50% and 90% credible intervals on d.

We found qualitatively similar results using a diversity metric related to Simpson’s concentration (which we refer to as diversity; *28*) and the time series of species’ relative abundances, though there were more cases where *Δ*γ > *Δ*α (Fig. 3b; Fig. S6). Species richness is more strongly influenced by rare species than our diversity measure (*28*), suggesting that our overall results are not primarily driven by rare species. Indeed, variation in the diversity measure used here shows that altered numbers of relatively abundant species, either at the α-scale of local sites or the larger γ-scale, can also drive changes in spatial β-diversity (Fig. 3b, Fig. S6). Numerically abundant species are often expected to contribute disproportionately to the provision of ecosystem function or services (*29–32*). Our results showing the importance of relatively abundant (or common) species for homogenization and differentiation, combined with their effects on ecosystem services, suggest that there is an urgent need to study the biodiversity dynamics of common species, not just the rare species that are often the focus of conservation biology.

Although we find some overall tendencies across all of our compiled data (i.e., local gains in species richness slightly outpace regional gains, leading to homogenization, especially for long duration checklist studies), our results suggest that changes in *α*-, *β*-, and *γ*-diversity through time are highly variable across locations, taxa, and time. This matches general findings on local diversity (*4–6, 33*), and population trends (*34, 35*), where variation in the direction of change means that the strength of overall net trends are weak and often statistically indistinguishable from zero. We argue that perhaps these results should not be surprising. Certainly humans are having many impacts that could lead to biotic homogenization, including transporting species beyond their historical biogeographic boundaries, and recreating urban or high intensity agricultural landscapes repeatedly (*12–14*). However, at the same time, many other impacts could lead to differentiation, including: substantial fragmentation of the landscape, the creation of strong spatial gradients of human impact intensity, applying spatially varying resource management practices and land use regulations, and climate change, which induces species to shift at different rates, all leading to spatial heterogeneity (*15, 16, 36*). In this context, we stress that the lack of strong overall scale-dependent biodiversity change and prevalence of homogenization should not in any way be taken to indicate that humans are not having a large impact on biodiversity. Both reductions in *β*-diversity, as in homogenization, or increases, as in differentiation, are likely a result of humans modifying nature. Furthermore, our focus on scale-explicit diversity, and temporal changes of spatial variation in species composition (i.e., the number of distinct assemblages within a region), is not able to track species identities. On top of the *α*-, *β* - and *γ*-diversity change shown here, for many of these assemblages the identity of species within them is also likely changing (*4, 6*). A loss of a shared species matched by a gain of a different shared species between communities has no net effect on *β* diversity but clearly involves change, possibly driven by humans. Simple descriptions for biodiversity change may often be hard to find.

Our results support moving beyond a belief in the predominance of ubiquitous biodiversity loss and homogenization, and instead working to understand the multiscale nature of biodiversity change and embracing its variability. Our conceptual classification emphasizes that different metacommunities are experiencing fundamentally different types of temporal change across scales, resulting in variable trends, and often no trend, in spatial *β*-diversity through time. Respectively, biotic homogenization and differentiation were often characterized by gains or losses in the number of regionally widespread, and/or relatively abundant species. This suggests that there is still time to protect many rare and narrow-ranged species before they ultimately contribute to homogenization. It also means that efforts to slow the spread of widespread (possibly non-native) species are vital for preventing biotic homogenization. Conversely, greater attention to variation and changes in the number of common species (*29, 37*), whether they be those with high occupancy within a region or having high relative abundance at either local or regional scales, will increase our understanding of both biotic homogenization and differentiation. Many species require protection across multiple sites or at landscape (or larger) spatial scales for effective conservation (*38, 39*). The simple conceptual typology introduced here simultaneously considers change in *α*-, *β*- and *γ*-diversity through time, showing how contemporary biodiversity monitoring for management and conservation can embrace a multiscale approach.

## Acknowledgments

SAB, TE, AS, RvK, WBX, JMC gratefully acknowledge the support of the German Centre of Integrative Biodiversity Research (iDiv) Halle-Jena-Leipzig (funded by the German Research Foundation; FZT 118). MD acknowledges funding by the European Union (CoralINT, GA 101044975). Views and opinions expressed are however those of the author(s) only and do not necessarily reflect those of the European Union or the European Research Council. Neither the European Union nor the granting authority can be held responsible for them. MD thanks Marten Winter for chocolate, and the Leverhulme Trust Research Centre – the Leverhulme Centre for Anthropocene Biodiversity. TE was supported by the German Research Foundation (DFG) within the project “Establishment of the National Research Data Infrastructure (NFDI)” in the consortium NFDI4Biodiversity (project number 442032008). AFE acknowledges the Fisheries Society of the British Isles Studentship. ISM received funding from the European Union Horizon 2020 research and innovation programme under the Marie Sklodowska-Curie grant agreement no. 894644. BJM acknowledges support from USDA Hatch grant MAFES #1011538 and NSF EPSCOR Track II grant #2019470.

## Funding

German Research Foundation; FZT 118; SAB, TE, AS, RvK, WBX, JMC. ERC GA 101044975 and the Leverhulme Centre for Anthropocene Biodiversity: MD

## Author contributions

**Conceptualization:** SAB, BM, TE, ISM, DM, HS, NJG, AM, MD, JMC

**Methodology:** SAB, BM, VB, CFYC, TE, AFE, ISM, DM, FM, AS, HS, RvK, WBX, NJG, AM, MD, JMC

**Investigation:** SAB, BM, VB, CFYC, TE, AFE, ISM, DM, FM, AS, HS, RvK, WBX, NJG, AM, MD, JMC

**Visualization:** SAB, BM, TE, ISM, DM, HS, NJG, AM, MD, JMC

**Writing – original draft:** SAB, BM, MD, JMC

**Writing – review & editing:** SAB, BM, VB, CFYC, TE, AFE, ISM, DM, FM, AS, HS, RvK, WBX, NJG, AM, MD, JMC

## Competing interests

The authors declare that they have no competing interests.

## Data and materials availability

All data used in this study are open access. R code used for data compilation (https://github.com/chase-lab/metacommunity_surveys, https://github.com/chase-lab/checklist_change, https://github.com/chase-lab/homogenisation-richness) and all analyses (https://github.com/sablowes/WhittakerBetaChange) are available, and will be archived in Zenodo prior to publication.

## Supplementary Materials

### Materials and Methods

#### Data compilation

Our conceptual typology requires estimates of species richness changes at two scales. We refer to them as local (*α*) and regional (*γ*), the exact definition of which varies among data sources. To make our data search and synthesis as comprehensive as possible, we searched broadly for data that met these criteria, where regions had at least four plots or locations, and where richness changes were estimated over a period of at least ten years. We started by identifying 80 relevant datasets within the BioTIME database (*20*) that monitored patterns of species abundances within assemblages. To this, we added: (1) similar assemblage-level time series of studies not (yet) included in BioTIME (e.g., *21, 22*); (2) data from studies using ‘resurveys’, where sites associated with a historical dataset were revisited and re-surveyed using similar methodology in more recent times; (3) data from ‘checklist’ studies where species known to be present in a given locality (and region) at a ‘historical’ point in time were indicated together with species present in that locality at a later point in time (minus those that went extinct from a site plus those that newly colonized that site); and, (4) data from studies that reported changes in species richness at two spatial scales, but for which the underlying raw data were not available. Because of the relatively specific data requirements, literature searches were conducted in an ad-hoc fashion, rather than using a formal literature search. In all, we compiled a total of 337 regions and a total of 20,470 locations that met our criteria; 283 regions documented repeated samples of species assemblages through time; 54 regions were compiled from checklist studies (Fig. S4; *40-311*).

#### Data standardization

To quantify changes in *β*-diversity that emerged from combined changes occurring at the local- and regional-scale, we required that the starting and end years for all locations within a given region were the same. This ensured that change estimated across the different locations within a region covered the same period of time, and meant that regional changes estimated by aggregating all species across all locations within regions also covered the same time period. Additionally, to ensure that our analyses did not quantify changes in species richness due to variation in sampling effort, we needed to standardize sampling effort (e.g., the number of plots or transects) across all locations for each time point within regions. The heterogeneous nature of the data that we compiled meant that we needed slightly different procedures to identify combinations of locations and years for different data sources. For clarity, we delineate broad categories of data structures, and describe separately how locations and years were selected and sample-effort standardization needed for the different structures.

#### Checklist data

Checklist data typically consisted of species lists for locations within regions, compiled from a historical time period and from a more contemporary time period (mean = 273 years; median = 258 years; range = 10 - 501 years). These lists were compiled either from samples and/or observations collected during the two periods, or more frequently, by counting native species only to determine the richness of the historical period, with the contemporary species richness calculated as the sum of native and introduced species (minus any species that went extinct). For our analyses, we selected regions that had at least four locations, removed locations that documented species lists for only one period, and finally, ensured that all locations within each region had data from the same time period for both the historical and contemporary species lists.

#### Resurvey data

We distinguish three different data structures that we refer collectively to as resurvey data:

i. Compiled time series data that document repeated samples of assemblages documenting species abundances, i.e., data from the BioTIME (20), RivFishTime (21), and InsectChange (22), plus similarly structured data that we compiled for this and related studies. We first filtered data to ensure that samples from all locations within regions had a temporal duration of at least ten years, and that a minimum of four locations within a region were sampled per year (allowing us to track diversity of the same locations through time within regions). Locations within regions were identified using geographic coordinates in the data, although we also included regions with only one geographic coordinate where discrete, unique samples could be identified (e.g., multiple plots within a site). After applying these filters, the number of locations sampled per year often varied considerably within regions, and we sought to identify locations, as well as start and end years that balanced a trade-off between the number of locations and the duration of the sampling period for each region. To do this, we first identified all year-pairs -- combinations of start and end year with at least ten years separating them -- for all locations within a given region. We then determined different thresholds for what proportion of the total number of locations we wanted to retain, using a combination of the total number of locations in a region, and visual inspection of locations sampled in each year. For example, for resurvey data newly collated for this study, we selected starting and end years where the proportion of the maximum locations was at least 90% for regions with fewer than 20 locations, 50% for regions with more than 20 locations, and 25% for the NERC Countryside survey data, which had between 60 and 300 locations across the UK (and where the lower threshold meant that the temporal duration of the surveys increased by more than ten years). For regions in the BioTIME and RivFishTime databases, we identified year-pairs with at least 75% and 90% of the maximum number of locations, respectively. Multiple year-pairs often remained following this, and we selected the pair of years with the longest duration, and finally, broke any remaining ties by selecting the pair of years with the most locations. For mosquito data sourced from Vectorbase (https://vectorbase.org/vectorbase/app) and compiled in InsectChange there were frequently fewer sites sampled monthly, and we visually selected locations and the start and end years for each region to ensure that we would be able to standardize the sampling of the same months for each location through time. Next, we ensured that sampling effort was consistent across all years and locations within regions, using sample-based rarefaction (*312*) where required to standardize effort. Note that for many data (e.g., data from InsectChange and other invertebrate data) where sampling took place across multiple months within years, we used sample-based rarefaction to resample equal numbers of samples across the same months for all locations within a region, which were then compiled to provide one sample per year for each location. Additionally, for data collected using multiple sampling methodologies (e.g., mosquitoes sampled using different attractants, or freshwater fishes collected with different techniques), we identified the methodology that ensured the maximum number of time series, and standardized sampling effort using data collected with only one method.
ii. We collated data from studies where sites associated with a historical dataset were revisited and re-surveyed using the same methodology in more recent times, sometimes referred to as “legacy” studies (e.g., *313*). Again, we required each region to have at least four locations and ten years or more between the historical and contemporary samples.
iii. Finally, we collated studies that estimated species richness changes at two scales, where there were at least four sites at the smaller scale and ten years between the first and last sample. For these studies raw data were not available (*n* = 15), but available data provided an estimate of the average local richness at two time points, and a single value for regional richness at two time points.

#### Estimating richness, diversity and its change

For the majority of the data, we calculated species richness from the effort-standardized locations and years as the number of distinct species, though higher classifications, such as genera, were sometimes used where studies only classified organisms to genus. We calculated species richness for each location within each region for every available year to document changes in α-scale species richness. γ-scale richness was calculated as the number of species in all sites combined for each region and each year. This method of calculating regional richness yields a single number for each region at each time point, and we calculated two types of resamples for regional richness. For datasets where sample-based rarefaction was not required to standardize effort, jackknife resamples were calculated by systematically leaving each location out of the regional richness calculation once. For where effort-standardization was more complex and required the use of sample-based rarefaction, we used 200 bootstrap resamples (i.e., richness resamples were estimated using all locations, not *n*_locations_-1); and then, to prevent these resampled data dominating the data to which models were fit, we subsampled the bootstrap resamples down to the same size as a jackknife would have been (i.e., we used a random subset of the bootstrap resamples equal to *n*_locations_ – 1 for the given dataset). Finally, we summarized both the jackknife and bootstrap resamples by calculating the median regional richness, to which models were fit.

Many data sources, including the 54 datasets with checklist data and 118 regions in the resurvey data, had only two years of data available (e.g., a historical and more contemporary sample). As a result, to maximize the number of regions in our complete analysis, we calculated richness change using the log-ratio of species richness in the most recent time point and species richness in the initial sample, divided by the number of years between the two samples (i.e., 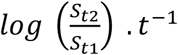, where *S*_t2_ is species richness in the most recent sample (t2 = year), *S*_t1_ is species richness in the first sample (t1 = year), and *t* = t2 – t1 + 1 is the number of years between the samples. This was done separately for each location in each region. These same data were aggregated and used to calculate concomitant changes in regional diversity through time, quantified as the log-ratio of resamples (either jackknife or bootstrap) of species richness at the regional scale in the most recent sample and the resample of species richness in the initial sample, divided by the number of years between the two samples.

In addition, because many of our data sources included both information on the abundance of individual species, as well as time series of more than two years, we also calculated diversity metrics that differ in their sensitivity to common and rare species (*28, 314*) for all years having effort-standardized data, and estimated rates of change using statistical models. Specifically, we calculated richness and the inverse of Simpson’s concentration (*28*). These two metrics are equal to diversity with order *q* = {0, 2}, where increasing *q* decreases the influence of rare species, and 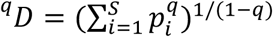; *p*_i_ is the frequency (relative abundance) of species *i* (*27*). These diversity measures are sometimes referred to as Hill numbers, numbers equivalents, or effective numbers (27, *314*), here we refer to them as species richness and diversity for simplicity.

#### Statistical models

To estimate α- and γ-scale richness changes, we fit multilevel (also called mixed effects or hierarchical) models to data from each scale separately. For our initial analysis of the changes calculated using log-ratios (*n*_regions_ = 337), these models took the form:

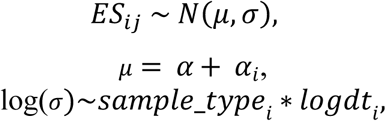

where *ES*_ij_ is assumed to have a Gaussian error distribution and is the *j*th local-scale estimate of species richness change in region *i*, or at the regional-scale the *j* subscript is dropped and *ES*_i_ is the median of the regional richness resamples for region *i; α* is the overall intercept and equal to the average rate of change estimated for each scale, and *α*_i_ is the departure from the overall intercept for each region (i.e., the varying intercept for regions). Because variation in the log-ratio estimates of change was a decreasing function of the duration between samples, we included covariates for residual variation (σ), where *sample_type*_i_ estimated a separate intercept according to whether region *i* were resurvey or checklist data, and *logdt*_i_ was the natural logarithm of the number of years between the first and last samples in region *i.* We fit models using Bayesian methods and we assumed the following, weakly regularizing priors:

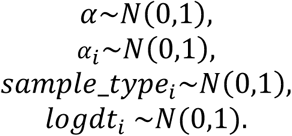

For the time series data of species richness and diversity (i.e., *^q^D*, {*q* = 0, 2}; *n*_regions_ = 165), we estimated rates of change using statistical models (rather than change being calculated using log-ratios as was done in the initial analysis). Time series data of diversity metrics were constrained to be positive values, but due to effort standardization were not always integer values, so we fit models that assumed lognormal error distributions and identity link functions. Overall rates of change were estimated by including (mean centered) year as a predictor, and we allowed rates of change to vary among regions by including varying slopes (for year) and intercepts for regions in the regional scale model, and both regions and locations within regions for the local scale model.

We fit all models using the Hamiltonian Monte Carlo (HMC) sampler Stan (*315*), and coded using the brms package (*316*). We fit models with variable numbers of chains and iterations to ensure posterior distributions had sufficient effective sample sizes. Visual inspection of the HMC chains and Rhat summaries showed model convergence (all Rhats < 1.05).

We used models fit to the *α*- and γ-scale data (described above) to quantify changes in *β-*diversity (Δ*β*). Estimates of Δ*α* and Δ*γ* were used as x- and y-coordinates, respectively, and then the distance of these points from the 1:1 line was calculated (Fig. 1). Specifically, we extracted 1000 draws from the posterior distribution of the *α*-scale estimate and designated them as the x-coordinate, and combined this with 1000 draws from the corresponding γ-scale estimate as the y-coordinate, and then calculated the distance from the 1:1 line for each point. This means the units of our estimate of *β*-diversity is effective numbers of communities (*27*); and as rates of change in the x-and y-values are on a log-scale, Δ*β* is also describes proportional changes. Variation among regions in Δ*β* were calculated similarly: estimates of change for individual regions used the varying intercepts (initial analysis of log-ratios) or varying slopes (time series data) from models fit to data at the α- and γ-scales.

## Supplementary Figures

**Figure S1:**
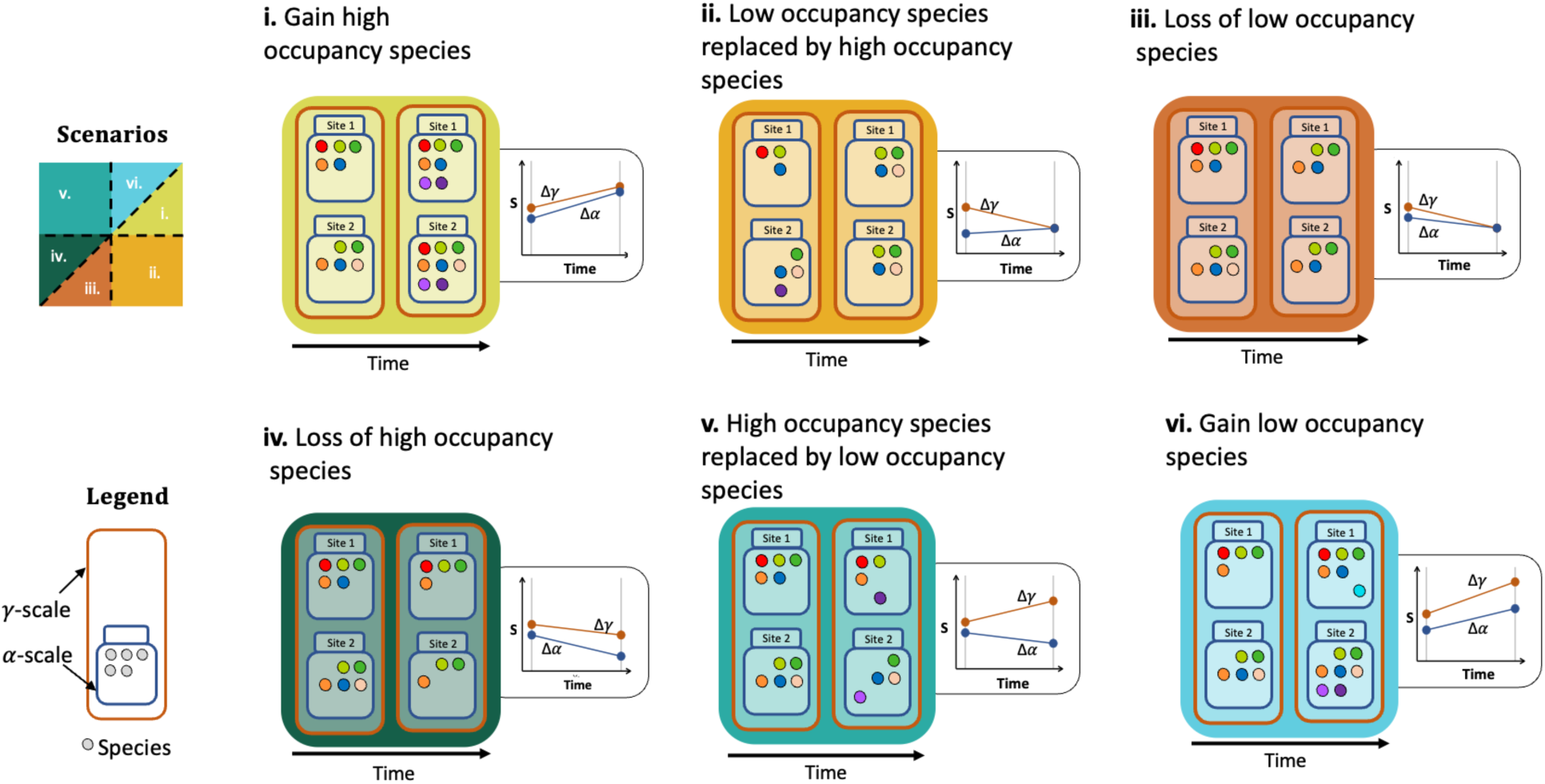
Simplified illustrations of species occupancy and richness changes underpinning each of the scenarios in. Figure 1. Each panel shows a simplified scenario of richness changes at smaller (*α*) and larger (*γ*) scales between two time points, accompanied by a regression showing richness at the two scales as a function time (slopes are labeled Δ*α* and Δ*γ* for each scale).

**Figure S2:**
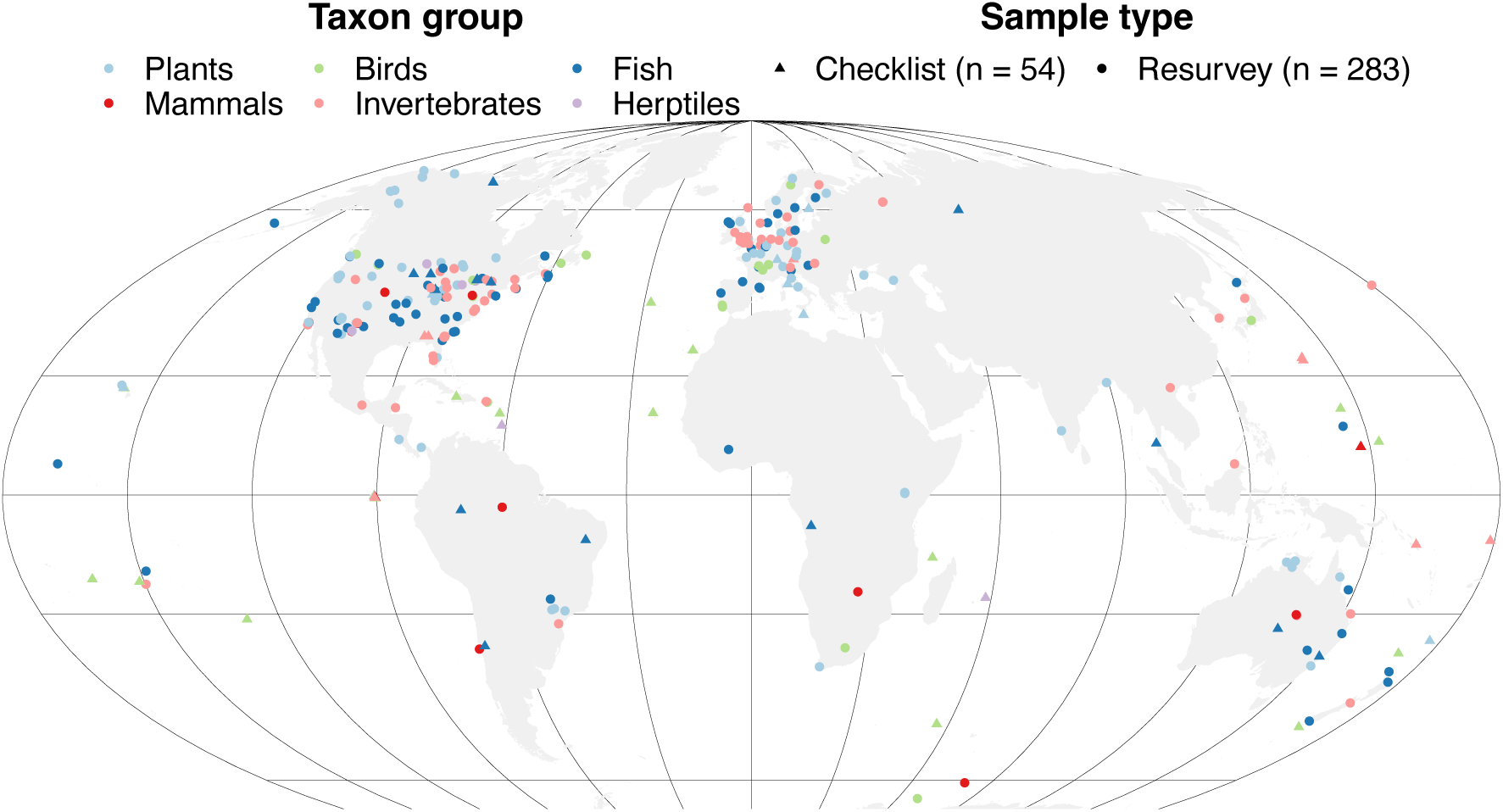
Map showing the central coordinate of each dataset (region) in our compiled data (n_regions_ = 337).

**Figure S3:**
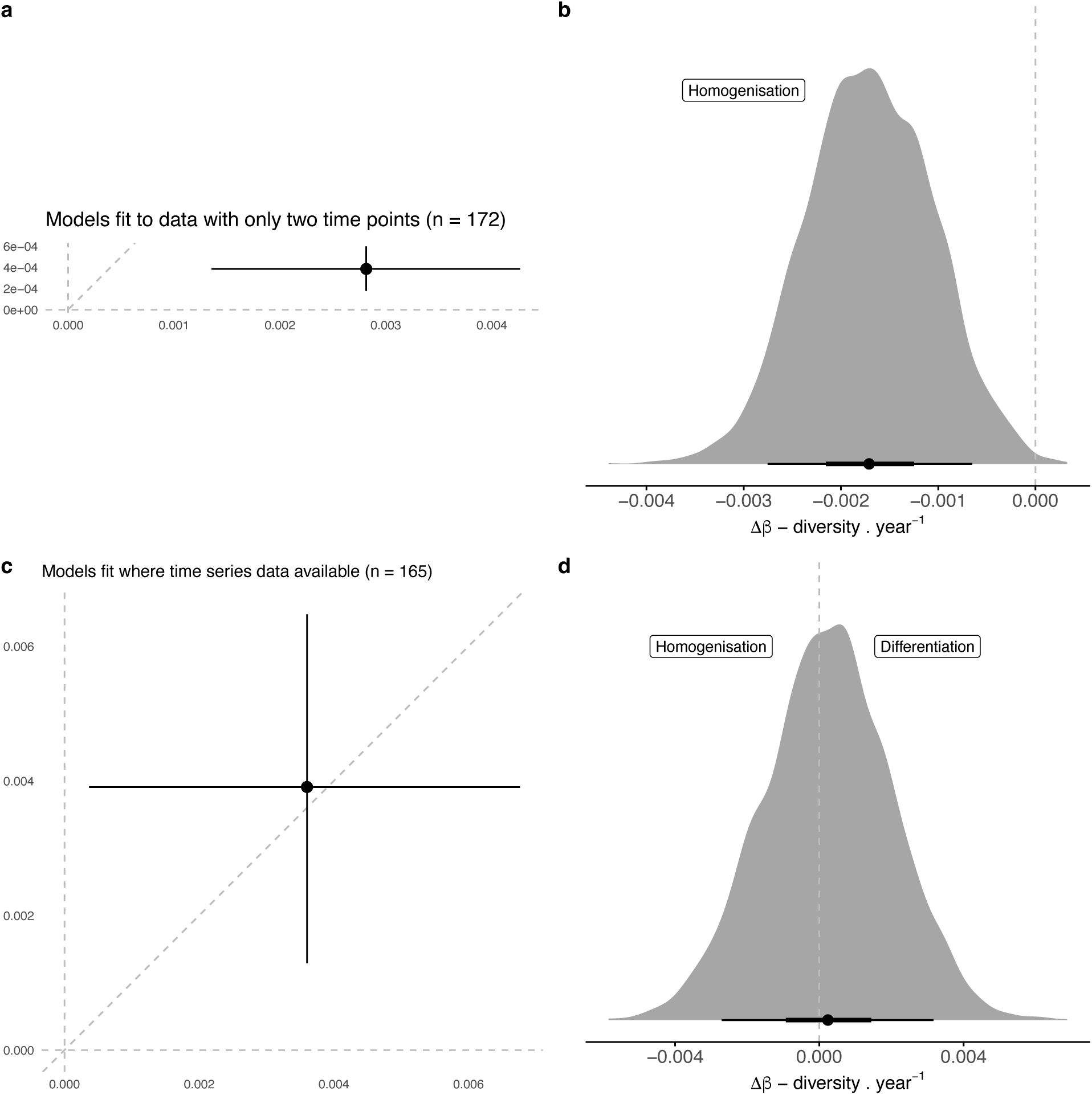
The tendency towards biotic homogenization is strongest in data where only two time points were available. (a) Change in γ-diversity as a function of change in α-diversity, and (b) change in beta-diversity estimated by models fit to data with only two time points; (c) change in γ-diversity as a function of change in α-diversity, and (d) change in beta-diversity estimated by models fit to data where time series were available. *Δ*β was calculated as the distance from 1:1 line (left = homogenization, right = differentiation) of 1000 draws of *α*- and *γ*-scale intercept posterior distributions; black point shows median, bar represents 50% (thick) and 90% (thin) credible intervals. Note all model results shown here were fit to duration standardized log-ratios as per Fig. 2 in the main text.

**Figure S4:**
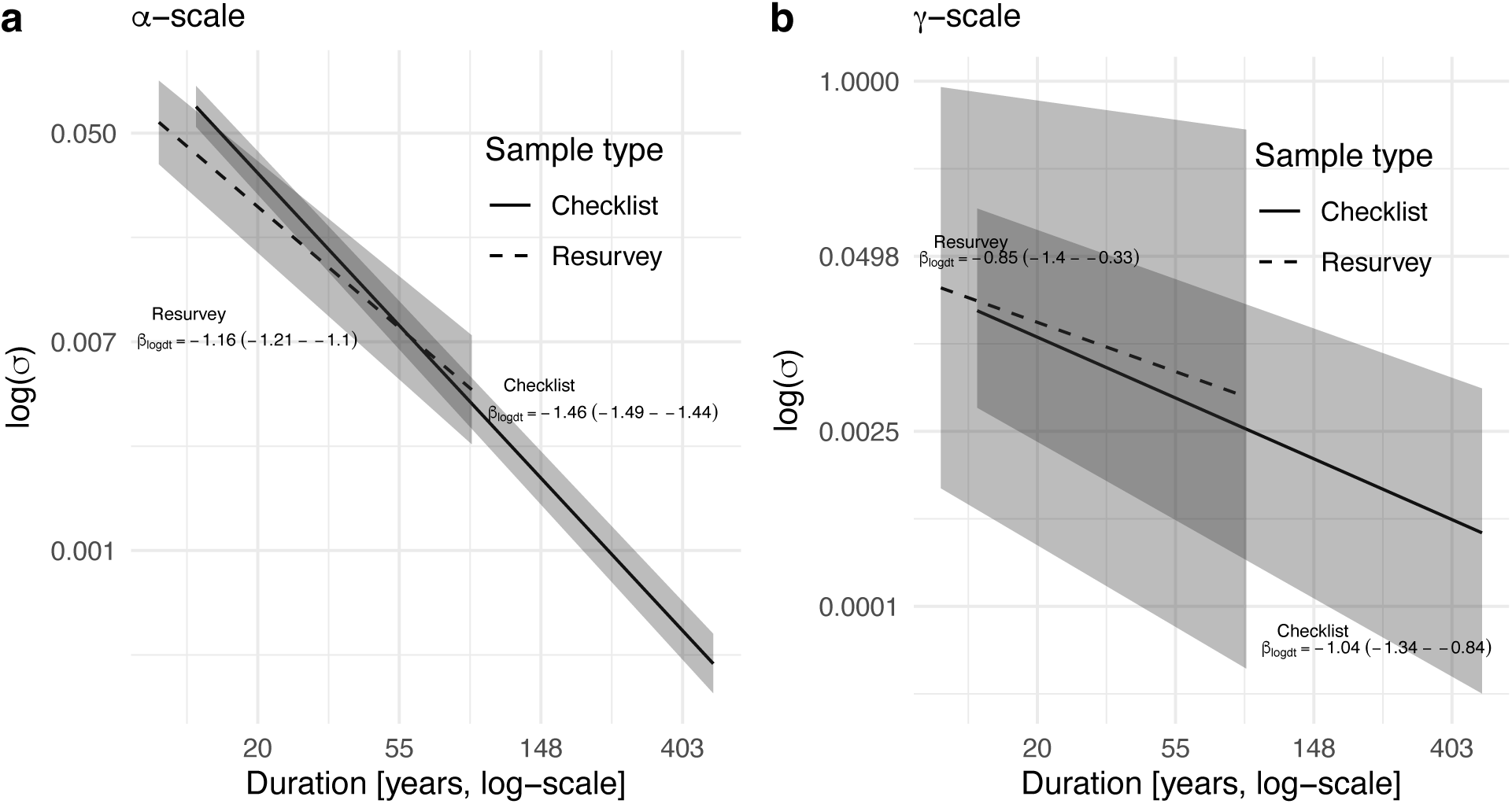
Scale-dependent residual variation as a function of temporal duration for the two sample types. Models fit to the full data set adjusted for residual variation being a decreasing function of temporal duration for both types of samples (checklists and resurveys) at the (a) α- and (b) γ-scales.

**Figure S5:**
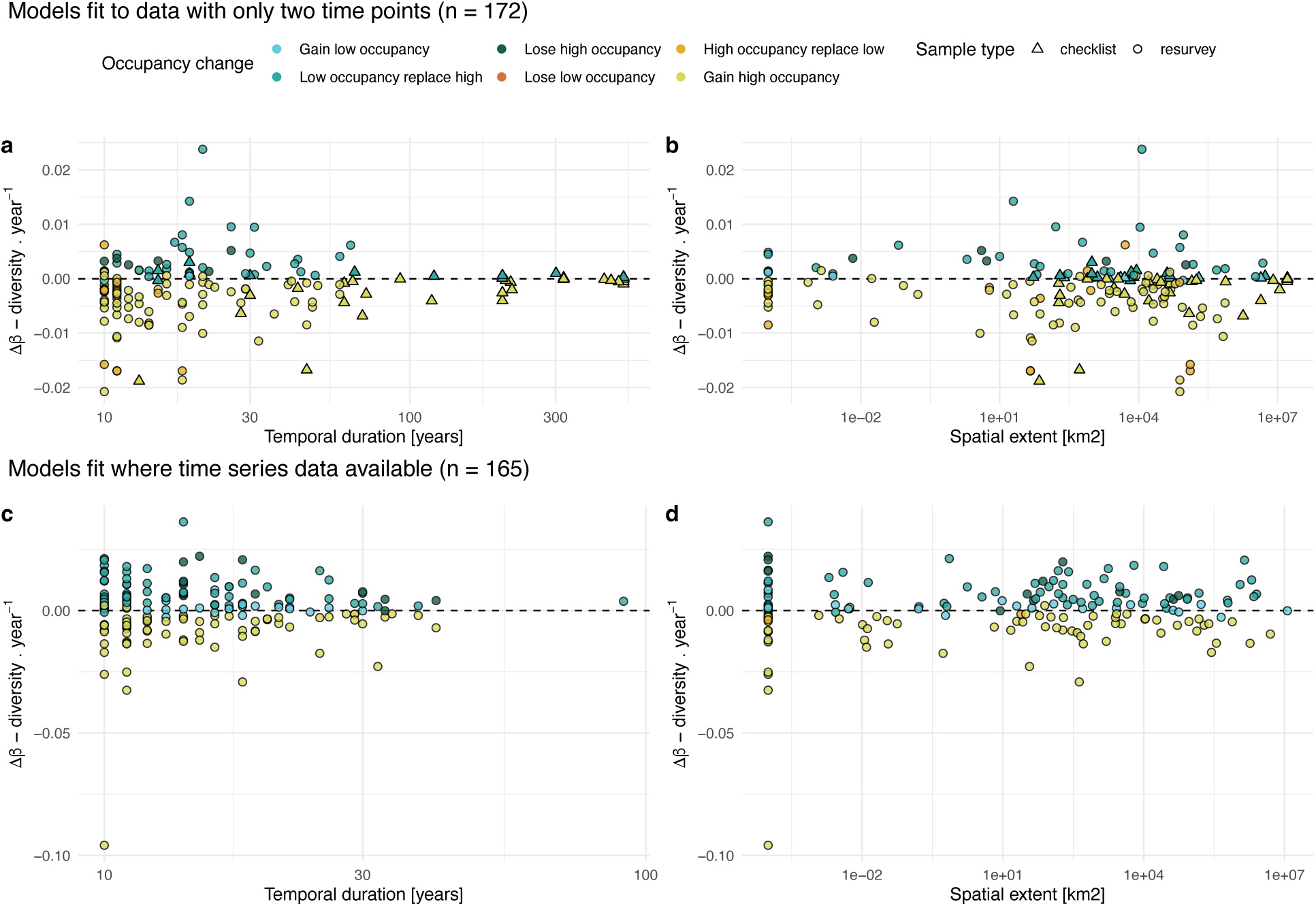
Biotic homogenization is strongly associated with data sets that had only two time points, and more common at intermediate to large temporal and spatial scales. Changes in beta-diversity as a function of (a) temporal duration and (b) spatial extent estimated using models fit to data with only two time points available; and, changes in beta-diversity as a function of (c) temporal duration and (d) spatial extent estimated using models fit to data where time series were available. Each point shows Δ*β* for an individual dataset (region) calculated as the mean distance from the 1:1 line (< 0 = homogenization, > 0 = differentiation) of 1000 draws of each *α*- and *γ*-scale regional estimate (i.e., overall intercept plus regional random intercept). Both x-axes are on a logarithmic scale.

**Figure S6:**
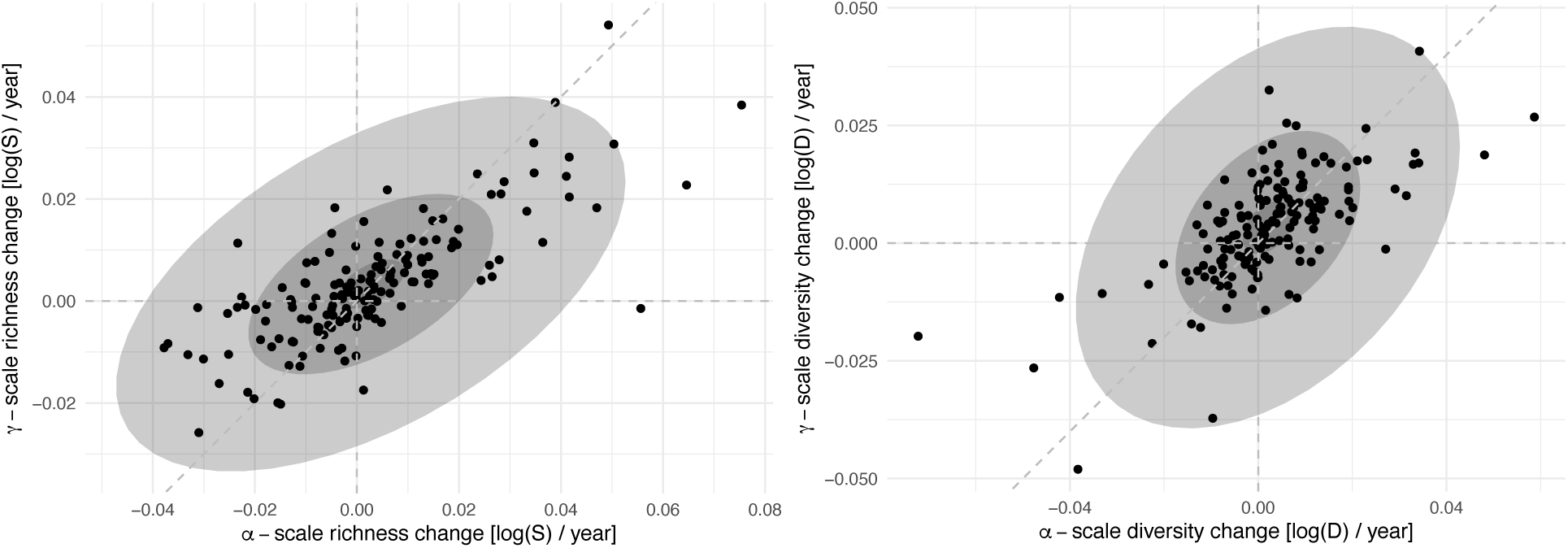
Scale-dependent variation in diversity change in the subset of datasets with species abundance time series. Changes in *α*- and *γ*-scale diversity of (a) species richness, and (b) diversity (the inverse of Simpson’s concentration (28), and more influenced by numerically abundant species). Shading on each panel shows the 50% and 90% quantiles of variation in the posterior distribution describing the varying rates of changes among the datasets from multi-level models fit to the *α*- and *γ*-scales separately. Each point represents a dataset (n = 165) and shows the median of the posterior of the *γ*-scale as a function of the *α*-scale. In (a), for richness, greater variation at the *α*-scale (relative to the *γ*-scale), shows how local species richness gains outpacing regional richness gains frequently results in homogenization, and richness losses (i.e., Δ*α* and Δ*γ* < 0) are more often associated with differentiation (i.e., Δ*γ* > Δ*α*) than homogenization. Variation in *α*- and *γ*-scale changes are more balanced among the datasets for diversity (b). The greatest rates of species gains (q = 0) were found for datasets at the *α*-scale. In contrast, the greatest rates of diversity loss were found for datasets at the *α*-scale for q = 2.

## Notes

### Competing Interest Statement

The authors have declared no competing interest.

### Summary of Updates

Revision improves initial analysis, and adds analyses of species relative abundance time series data.

